# Vaccination against helminth IL-33-modulators permits immune-mediated parasite ejection

**DOI:** 10.1101/2024.08.22.609138

**Authors:** Danielle J Smyth, Suzanne Hodge, Nicole Ong, Josh Richards, Florent Colomb, Vivien Shek, Tania Frangova, Henry J McSorley

## Abstract

The murine intestinal nematode *Heligmosomoides polygyrus bakeri* powerfully modulates the host immune response. This is achieved in part through the HpARI family (HpARI1/2/3), which act on IL-33, and the HpBARI family (HpBARI and HpBARI_Hom2), which act on ST2. Here, we find that this immunomodulation is evident only in the first week of infection, with abrogation of ST2 detection and systemic suppression of IL-33-dependent responses. Vaccination with individual HpARI or HpBARI family members raised antibody responses which could block these proteins’ immunomodulatory activities. During infection, vaccination could release the host from immunosuppression: HpARI2 vaccination resulted in much increased ILC2 and Th2 immunity, with heightened serum IL-4 and IL-5 responses, but did not abrogate ST2 suppression. In contrast, a HpBARI+HpBARI_Hom2 vaccination cocktail resulted in abrogation of ST2 suppression, and again increased Th2 immunity and serum cytokine responses. Either of the HpARI2 or the HpBARI cocktail vaccinations provided significant protection against subsequent *H. polygyrus bakeri* infection. We therefore show a proof of principle that vaccination with immunomodulatory proteins can protect the host against infection, and can be used as a tool for blocking the effects of specific parasite-derived proteins.

## Introduction

Intestinal nematode infections affect over 1 billion people worldwide. These parasites can cause significant morbidity (especially in children), including growth stunting, poor school performance, anaemia and malnutrition ^1^. Veterinary parasites of livestock are also of large economic importance, and it is estimated that in Europe €1.8 billion per year is lost to intestinal nematode infections ^2^. Although anthelmintic agents are available for these pathogens, drug resistance is becoming common in livestock parasites, with evidence of development in human-infective nematodes ^3^. Furthermore, drug treatments must be given regularly in endemic areas, with rapid reinfection after drug clearance. Therefore, an effective vaccine which provided long-term resistance to helminth infection would be highly advantageous. Currently, there are no vaccines available for any human-infective nematodes, and only very limited vaccines are available against nematodes of veterinary importance ^4^. Animal models of helminth infection such as *Heligmosomoides polygyrus bakeri* (*Hpb*) are useful in investigation of anti-parasite immunity and the development of effective anti-parasite vaccines ^5, 6^. *Hpb* is a widely-used model parasite: it is an intestinal nematode of mice, and forms chronic infections associated with immunomodulation of bystander responses ^7^.

Recently, multiple immunomodulatory proteins have been identified in *Hpb* secretions. They contain multiple unique immunomodulatory protein families, including the HpARIs, the HpBARIs, and the Hp-TGMs, as well as many secreted protein families shared with other parasitic nematodes, such as the apyrases, the DNAses, and the VAL family ^8^ ^9^. Of note, both the HpARIs and the HpBARIs act on the same immunological pathway – the IL-33 response. HpARI1 and HpARI2 act directly on IL-33, binding to the cytokine, preventing interaction with its receptor ^10, 11, 12^. HpARI1 and HpARI2 also bind to DNA and the extracellular matrix, which extends their half-life in vivo and allows them to tether IL-33, preventing its release ^13^. HpARI3 also binds to IL-33 but stabilises the cytokine, amplifying its effects in vivo ^13, 14^. The HpBARI family, conversely, act on the IL-33 receptor, ST2. Both HpBARI and HpBARI_Hom2 bind with high affinity to the IL-33 receptor, preventing its interaction with IL-33. We previously showed that HpBARI binding to ST2 was sufficient to abrogate detection of ST2 on the cell surface using commercially available antibodies, thus the disappearance of the ST2 signal by flow cytometry can be used as a proxy for the HpBARI family’s effects ^15^. Our previous work showed that both HpARI2 and HpBARI were capable of suppressing responses in Alternaria-induced, IL-33-dependent models of asthma ^10, 15^, however due to a lack of transgenesis methodologies in *Hpb* ^16^, we could not define a role for these proteins during infection.

IL-33 is a pleiotropic cytokine, and depending on context IL-33 release can lead to allergy, anti-helminth responses, beiging of adipose tissue, anti-viral responses or immunosuppression ^17^. The IL-33 receptor consists of ST2 and IL1RAcP, and is expressed on mast cells, ILC2, Th2 cells, regulatory T cells, and activated Th1, CD8+ CTL and NK cells ^18^. IL-33 receptor signalling on mast cells results in neutrophil recruitment ^19^, while signalling on ILC2 and Th2 cells results in the release of IL-5 and eosinophil recruitment ^20^. Some populations of regulatory T cells (especially in the adipose tissue and colon) activate and expand in response to IL-33 ^21^. On Th1, CD8+ CTLs and NK cells, IL-12 treatment upregulates the IL-33 receptor, and subsequent IL-33 signalling is a potent signal for IFN-γ release and is important for resistance to viruses ^22^. Therefore the site and context of release is critical to the response to the cytokine.

In this study, we demonstrate that members of both the HpARI and HpBARI immunomodulatory protein families provide protection against infection in a vaccination regime. We furthermore show that vaccination releases the immune system from immunomodulation, through raising blocking antibody responses. These findings provide a mechanistic underpinning and proof of principle that immunomodulatory proteins are good candidates for vaccines against parasitic helminths.

## Materials and Methods

### Protein production

As described previously, HpARI1, HpARI2, HpARI3 proteins were expressed recombinantly with C-terminal myc and 6His tags ^14^, while HpBARI and HpBARI_Hom2 were expressed recombinantly with N-terminal 6His and myc tags ^15^. The HPOL_0001072601 (control protein) sequence is available at Wormbase ParaSite ^23^ and was expressed recombinantly with an N-terminal 6His tag. All sequences were cloned into a pSecTAG2A vector backbone (Thermo Fisher Scientific), and transfected into mammalian Expi293F cells using the Expifectamine transfection kit (Thermo Fisher Scientific). Proteins were purified from cell-free supernatants 7 days after transfection using Ni-NTA chromatography, and subsequently dialysed into 1 x PBS, filter sterilized and assayed for LPS levels using the HEK-Blue LPS Detection kit 2 (InvivoGen). All proteins used in these experiments contain < 0.1 EU/ug LPS.

### Animals

C57BL/6JCrl mice were purchased from Charles River, UK, while heterozygous IL-13^+/eGFP^ mice on a C57BL/6J background were provided by Prof Andrew McKenzie ^20^. Mouse accommodation and procedures were performed under UK Home Office licenses (project license PP9520011) with institutional oversight performed by qualified veterinarians. Mouse sex and age, as well as biological replicates per group in each experiment are stated in figure legends.

### Vaccination and Hpb infection

Mice were vaccinated with recombinant HpARI1, HpARI2, HpARI3, HpBARI, HpBARI_Hom2 or control protein (HPOL_0001072601), or PBS controls received PBS in adjuvant only. Mice were subcutaneously injected in the flank with 10 μg of parasite protein in 100 μl Alhydrogel (InvivoGen) alum adjuvant, followed by two boosts of 1 μg of parasite protein in adjuvant 28 and 35 days later. In HpBARI+HpBARI_Hom2 vaccinations, 10 μg (primary) or 1 μg (boost) of each protein were used in combination. At day 42 after the first injection, mice were infected with 200 *Hpb* L3 larvae by oral gavage. Mice were culled 7 days after infection for measurement of immune responses, or 28 days after infection for adult worm counts. Faecal pellets were collected at day 14, 21 and 28 of infection for egg counts.

In some experiments, mice were infected with *Hpb* without vaccination and culled at various timepoints as indicated in the figures. In other experiments, uninfected or *Hpb*-infected mice were intranasally administered 50 μg Alternaria allergen (Greer, XPM1D3A25) under isoflurane anaesthesia, and culled 24 h later.

### Tissue preparation for flow cytometry

Bronchoalveolar lavage (BAL): BAL was collected using a 26G needle inserted into the trachea of the mouse and 4 x 0.5 ml washes made using 1 x PBS. Cells were separated from the BAL fluid by centrifuging at 300 g for 5 min at 4 °C. Pelleted cells were used for flow cytometry, while supernatants were stored at -70 °C for cytokine ELISA.

Peritoneal lavage (PL): Peritoneal cells were collected by 2 x 5 ml lavages with ice-cold 1 x PBS in the peritoneal cavity. Cells were separated from the wash fluid by centrifugation at 300 g for 5 min.

Perigonadal White Adipose Tissue (pgWAT): Perigonadal adipose tissue pads were dissected, weighed and minced into small pieces using scissors. Tissue was placed into 1 ml digestion mixture comprising of 1 x PBS^++^ (PBS containing magnesium chloride and calcium chloride; Gibco) containing Liberase TM (50 µg/ml; Roche) and DNAse1 (160 U/ml; Sigma) and placed in a shaking incubator for 35 min at 200 rpm, 37 °C. Following digestion, 10 mM EDTA (Gibco) was added and cells were incubated without shaking for a further 5 min. Bijous were topped up with ice cold FACS buffer (1 x PBS, 0.5% BSA [Sigma]) and samples were filtered through a 70 µm filter (Greiner), and centrifuged at 300 g for 15 min at 4 °C. After centrifugation, the adipocyte layer was carefully removed, and the remaining stromal vascular fraction (SVF) was collected for flow cytometry staining.

Mesenteric Lymph Node: The mesenteric lymph nodes (MLN) were taken from each mouse and crushed through a 70 μm nylon filter (Greiner) then resuspended in complete RPMI media (RPMI-1640 (Gibco) supplemented with 10% FCS (Invitrogen), 100 U/ml penicillin, 100 μg/ml streptomycin (Gibco) and 2 mM L-glutamine (Gibco)) for counting and flow cytometry staining.

Lung: Either half (complete lobe) or three quarters (complete lobe plus largest segmented lobe) were taken from each mouse and collected in 900 μl 1x PBS^++^. Liberase TL (2 U/ml; Roche) and DNAse1 (160 U/ml; Sigma) were added to each collection tube and lungs were thoroughly minced with scissors. The minced lungs were incubated for 35 min, shaking at 200 rpm at 37 °C. The digest was then stopped with 5 ml ice cold complete RPMI (Gibco) and crushed through a 70 μm nylon filter (Greiner).

For all samples, red blood cells were lysed in ACK lysis buffer (Gibco), then resuspended in PBS for counting and flow cytometry staining.

### Flow cytometry

Single cell suspensions of cells were prepared as described as above before being resuspended in 1 x PBS containing a LIVE/DEAD fixable dye (Fixable Aqua Dead Cell Stain Kit (1:1,000, Invitrogen); or Zombie UV Fixable Viability Kit (1:200, BioLegend)) for 15 min in the dark at room temperature. Cells were centrifuged (400 g, 5 min, 4 °C) and resuspended in FACS buffer (1 x PBS supplemented with 0.5% BSA) containing purified anti mouse CD16/32 (Clone: 93, BioLegend, 1:50) and incubated at 4 °C for 20 min in the dark. Cells were then washed with FACS buffer, centrifuged as before and resuspended in FACS buffer containing extracellular antibodies and incubated for 30 min at 4 °C in the dark.

Extracellular antibodies used: CD127-FITC (clone: A7R34, 1:200); CD44-PerCP (clone: IM7, 1:200); CD90.2-AF700 (clone: 30-H12, 1:200); CD45-APCCy7 (clone: I3/2.3, 1:200), CD45-AF700 (clone: 30-F11, 1:200); CD3-Pacific Blue (clone: 17A2, 1:200), CD3-Biotin (clone: 145-2C11, 1:200), CD3-FITC (clone: 145-2C11, 1:200); CD5-Pacific Blue (clone: 53-7.3, 1:200); NK1-1-Pacific Blue (clone: PK136, 1:200); B220-Pacific Blue or -Biotin (clone: RA3-6B2, 1:200); CD11b-Pacific Blue, -FITC or -Biotin (clone: M1/70, 1:200); CD11c-Pacific Blue (clone: N418, 1:200); ST2-Biotin (clone: DIH9, 1:100); CD25-BrilliantViolet 650 (clone: PC61, 1:200); CD4-BrilliantViolet 711 (clone: GK1.5, 1:200), CD4-PE-Dazzle (clone: RM4-5, 1:200); KLRG1-PE-Dazzle or -PerCP (clone: 2F1/KLRG1, 1:200); Gr1-Biotin or -FITC (clone: RB6-8C5, 1:200); Ter-119-Biotin (clone: TER-119, 1:200); CD19-FITC (clone: 6D5, 1:200); FcεRI-PE-Cy7 (clone: MAR-1, 1:200); cKit-Pacific Blue or -PerCP (clone: 2B8, 1:200); ICOS-PE (clone: C398.4A, 1:200); IgE-PE (clone: RME-1, 1:200); Streptavidin-BrilliantViolet 510 or -PE (1:100), all from Biolegend; or CD5-FITC (clone: 53-7.3, 1:200); ST2-APC (clone: RMST2-2, 1:100); or CD49b – FITC (clone: DX5, 1:200) from Invitrogen.

If intracellular staining was not required, cells were washed with FACS buffer and centrifuged as before and resuspended in either FACS buffer or 1 x PBS for flow cytometry analysis. For cells undergoing intracellular staining, cells were washed with FACS buffer, centrifuged as before and resuspended in 250 µl in FoxP3 fixation/permeabilisation buffer (eBioscience) for 30 min at 4 °C in the dark. Cells were centrifuged as before and washed with 250 µl FoxP3 1 x permeabilisation buffer (eBioscience), centrifuged as before and resuspended in 1 x permeabilisation buffer containing intracellular antibodies.

Intracellular antibodies used: GATA3-PE (clone: TWAJ, 1:50); RORγt-APC (clone: B2D, 1:100); FoxP3-PE-Cy7 (clone: FJK-16s, 1:100), all from Invitrogen.

Cells were incubated for 30 min at 4 °C in the dark, washed with 1 x permeabilization buffer, centrifuged as before and resuspended in 1 x PBS or FACS buffer for flow cytometry analysis. Samples were acquired on an LSR Fortessa (BD) and analysed using Flowjo v10.9 (BD).

Gating strategies are shown for peritoneal lavage mast cells (live, CD45^+^ckit^+^lineage^−^FcεRI^+^) and ILC2 (live, CD45^+^KLRG1^+^lineage^−^) (**Supplementary Fig 1**); mesenteric lymph node ILC2 (live, CD45^+^lineage^−^CD127^+^GATA3^+^RORγt^−^) or Th2 cells (live, CD45^+^CD4^+^lineage^+^GATA3^+^Foxp3^−^) (**Supplementary Fig 2**); perigonadal white adipose tissue ILC2 (live, CD45^+^lineage^−^KLRG1^+^CD25^+^) (**Supplementary Fig 3**); and lung ILC2 (live, CD45^+^lineage^−^ICOS^+^CD90.2^+^CD25^+^CD4^−^FcεRI^−^) (**Supplementary Fig 4**). Lineage stains contained CD3, CD19, CD5, GR1, CD11b and CD49b.

### Serum antibody and cytokine measurements

Serum from each mouse was obtained by using injectable anaesthesia overdose followed by cutting of the brachial artery. Blood was collected into a serum Microtainer SST tube (BD), mixed and left for 30 min at RT before being centrifuged at 6,000 g for 3 min to separate the red blood cells from serum and was then subsequently frozen at -70°C until required for assays.

Antigen-specific antibodies in the serum were measured by ELISA, coating Nunc 96 well plates with 1 μg/ml of the individual HpARIs or HpBARIs in 0.1 M carbonate buffer pH 9.6. Serum was firstly diluted 1:500 with ELISA block (2% BSA in 1 x TBS / 0.05% Tween 20) and then serially diluted. Horseradish peroxidase-conjugated Goat anti-Mouse IgG (H+L) (BioRad) (1:2000) was used for detection, and was then developed with 1 x TMB (BioLegend) followed by a stop acid solution (1 M H_2_SO_4_). Plates were read at OD450 on a spectrophotometer (BMG Labtech ClarioStar). End-point titres were determined by identifying the reciprocal dilution of the mean plus 3 x SD of blank wells.

Cytokine measurements of serum were performed using LEGENDplex murine Th1/Th2 8 -plex Version 3 (BioLegend) as per kit instructions with data acquired using a Fortessa (BD) flow cytometer and analysed using software by BioLegend. The limit of detection for each cytokine is marked on the graphs as a dotted line.

Murine IL-5 cytokine measurements of BAL and IL-33 response assay supernatants in Supplementary Figure 5 were performed by ELISA using the mouse IL-5 uncoated ELISA kit (Invitrogen).

### IL-33 response assay

In vitro modulation of IL-33 responses was measured as described previously ^12, 14^. The CMT64 cell line (ECACC 10032301; RRID:CVCL_2406) was cultured in complete RPMI (as described previously) to confluency in 96-well plates (200 μl culture volume), in the presence or absence of HpARI1, HpARI2 or HpARI3 (100 ng/ml), and 2.5% serum from vaccinated animals as indicated. Cultures were immediately frozen on dry ice (causing cellular necrosis and release of stored IL-33), and stored at -70 °C. Cells were then thawed at 37 °C for 2 h and supernatants added directly to murine bone marrow cultures. 5 x 10^5^ cells/well of mouse bone marrow cells were cultured in the presence of 50 μl CMT64 freeze-thaw supernatants (containing HpARIs and serum as described above), IL-2 and IL-7 (10 ng/ml each (Biolegend)). Cells were cultured for 5 days, prior to collection of supernatants for IL-5 ELISA (Invitrogen).

### Statistics

Data was analysed using Prism version 10.2.3 (Graphpad). Statistical tests are detailed in figure legends: where multiple groups are compared one-way ANOVA with Dunnett’s multiple comparisons test is used. Where multiple groups and timepoints are compared, two-way ANOVA and Dunnett’s multiple comparisons test is used. Where two groups are compared, Mann-Whitney U test was used. All error bars indicate standard error of mean. Mouse numbers and sexes are indicated in figure legends.

## Results

### Hpb infection suppresses IL-33-dependent responses during the first week of infection

We previously showed that during *Hpb* infection, the action of the HpBARI family resulted in suppression of ST2 detection in vivo at day 7 of infection ^15^. To investigate how sustained and localised this ST2 suppression is, we carried out a timecourse of *Hpb* infection, using flow cytometry to assess surface ST2 on ILC2 and mast cells (constitutively ST2-positive cells) from the peritoneal cavity, mesenteric lymph node (MLN), perigonadal white adipose tissue or lung. We found that the profound suppression of ST2 detection was evident on both mast cells and ILC2s from all tissue sites assessed, however was restricted to the first week of infection only (**Fig 1A-E**). This indicates that the effects of the HpBARIs are systemic, but limited to the early phase of infection when *Hpb* resides within the tissue of the duodenal wall, and disappears when *Hpb* emerges into the lumen of the intestine, around day 10 ^24^.

**Figure 1:**
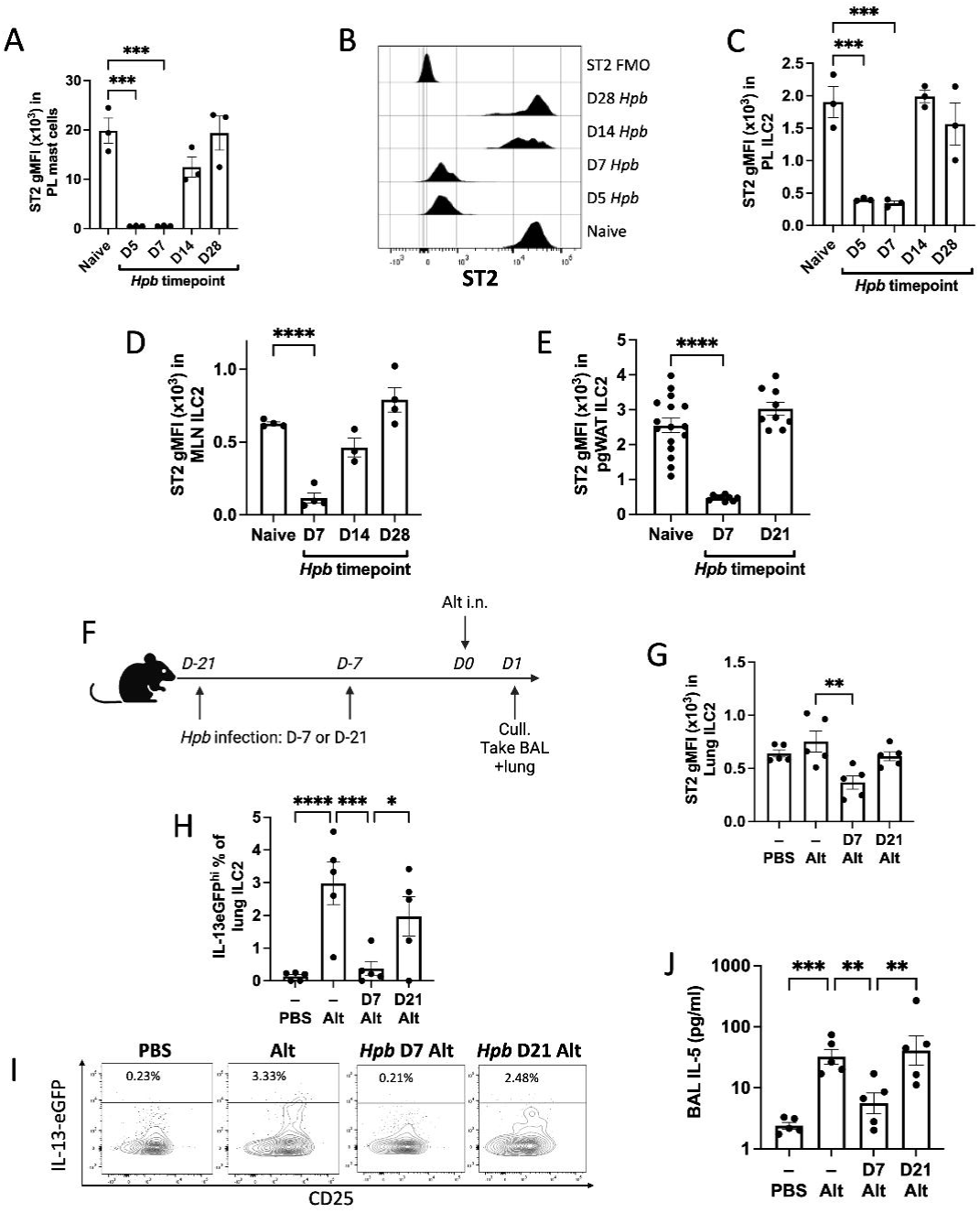
Systemic suppression of IL-33 responses during first week of *Hpb* infection. Mice were infected with *Hpb* and culled at day 5, 7 14, 21 and 28 of infection as indicated. ST2 geometric mean fluorescence intensity (gMFI) on peritoneal lavage (PL) CD45^+^Lin^−^ckit^+^FcERI^+^ mast cells (A), with representative ST2 histograms on gated mast cells (B). ST2 gMFI on PL CD45^+^Lin^−^KLRG1^+^ ILC2 (C), on mesenteric lymph node (MLN) CD45^+^Lin^−^CD127^+^CD90^+^RORgt^−^GATA3^+^ ILC2 (D) and on perigonadal white adipose tissue (pgWAT) CD45^+^Lin^−^CD25^+^KLRG1^+^ ILC2 (E). Experimental protocol for G-J: IL-13^+/eGFP^ mice were infected with *Hpb*, then either 7 days or 21 days later 50 μg Alternaria (Alt) allergen was administered intranasally (i.n) and mice were sacrificed 1 day later (F). ST2 gMFI on lung CD45^+^Lin^−^CD90.2^+^CD4^−^FcεR1^−^ ILC2 (G), IL-13eGFP^hi^ % of lung ILC2 (H), with representative IL-13eGFP versus CD25 bivariate plots of gated ILC2 (I). IL-5 concentration in bronchoalveolar lavage (BAL) (J). All data representative of at least 2 repeat experiments, using mice 8-15 weeks old. In A-D: n=3 per timepoint, male C57BL/6J mice. In E: n= 9-16 per group, both male and female C57BL/6J mice. In G-J: n=5 per group, male and female IL-13eGFP^hi^ mice. Analysed by one-way ANOVA followed by Dunnett’s multiple comparisons test. Unless otherwise stated, differences are not significant. * = p<0.05, ** = p<0.01, *** = p<0.001, **** = p<0.0001.

To determine if this systemic abrogation of ST2 detection was associated with functional suppression of IL-33 responses, we measured the IL-33-dependent response 24 h after intranasal Alternaria allergen administration. Alternaria allergen administration to the lung causes an acute and potent release of IL-33 due to protease-mediated damage and necrosis of epithelial cells, resulting in the release of ATP and IL-33. IL-5 and IL-13 production in response to Alternaria allergen are highly dependent on IL-33 signalling on lung ILC2 ^25, 26, 27^. In these experiments, we used IL-13^+/eGFP^ mice ^20^ to allow identification of ILC2 IL-13 production. Alternaria was administered to naïve IL-13^+/eGFP^ mice, or mice infected with *Hpb* 7 days or 21 days prior (**Fig 1F**). Similar to results seen in other tissue sites, we found that ST2 detection on lung ILC2 was suppressed 7 days after *Hpb* infection, and returned to baseline levels by day 21 of infection (**Fig 1G**). Alternaria allergen administration induced strong ILC2 IL-13 production in the lungs of uninfected mice, as well as IL-5 release to the bronchoalveolar lavage (**Fig 1H-J**). These responses were suppressed at day 7 of *Hpb* infection, but returned to similar levels as the positive control by day 21 of *Hpb* infection. Together, these data indicate that suppression of IL-33 responses is limited to the early phase of *Hpb* infection (when *Hpb* expression of HpARI and HpBARI families is highest ^14, 28^), and rapidly returns to baseline once the parasite enters the lumen of the intestine.

### HpARI2 (but not HpARI1 or HpARI3) vaccination abrogates immune suppression

To determine whether this suppression of IL-33 in the first week of *Hpb* infection was dependent on the effects of the HpARI and HpBARI families, we used a vaccination approach to raise a blocking antibody response against each immunomodulatory protein, focussing first on the HpARI family. In non-vaccinated mice, no antibody responses could be detected against any HpARI protein at day 7 of infection, and only became evident against specific proteins after day 14 of infection (**Fig 2A**). No detectable antibody response was raised against HpARI3 at any timepoint of *Hpb* infection, which may reflect its low expression level ^14^. HpARI1, HpARI2 or HpARI3 were administered in a vaccination regime with an alum adjuvant, followed by *Hpb* infection (**Fig 2B**). Serum antibody responses were assessed at day 7 of infection, when it was again seen that there was a negligible response against HpARI proteins in PBS-vaccinated controls. By contrast, HpARI-vaccinated mice raised a large antibody response against the vaccinated targets. Significant levels of cross-reactivity were seen between HpARIs, however antibody titres were in each case highest to the vaccinated HpARI family member (**Fig 2C**). Of note, HpARI vaccination produced antibody titres at least 10-fold higher than that seen at any timepoint in natural infection (**Fig 2A**). This antibody response could block the effects of the respective HpARIs in an in vitro IL-33 response assay, with anti-HpARI1 or HpARI2 serum tending to prevent the IL-33-blocking effect of HpARI1 and HpARI2 respectively, while anti-HpARI3 serum tended to reduce the known IL-33-amplifying effect of HpARI3 (**Supplementary Fig 5**). In vivo, HpARI2 vaccination increased Th2 (**Fig 2D**) and ILC2 responses (**Fig 2E**) in the mesenteric lymph node at day 7 of *Hpb* infection, compared to PBS-vaccinated infected controls. Serum IL-4 and IL-5 levels were also increased with HpARI2 vaccination (**Fig 2F, G)**, while IFN-γ was unchanged (**Fig 2H**). By each of these measures, HpARI1 had a similar but smaller effect than HpARI2, and trends for increases with HpARI1 did not reach statistical significance in these experiments.

**Figure 2:**
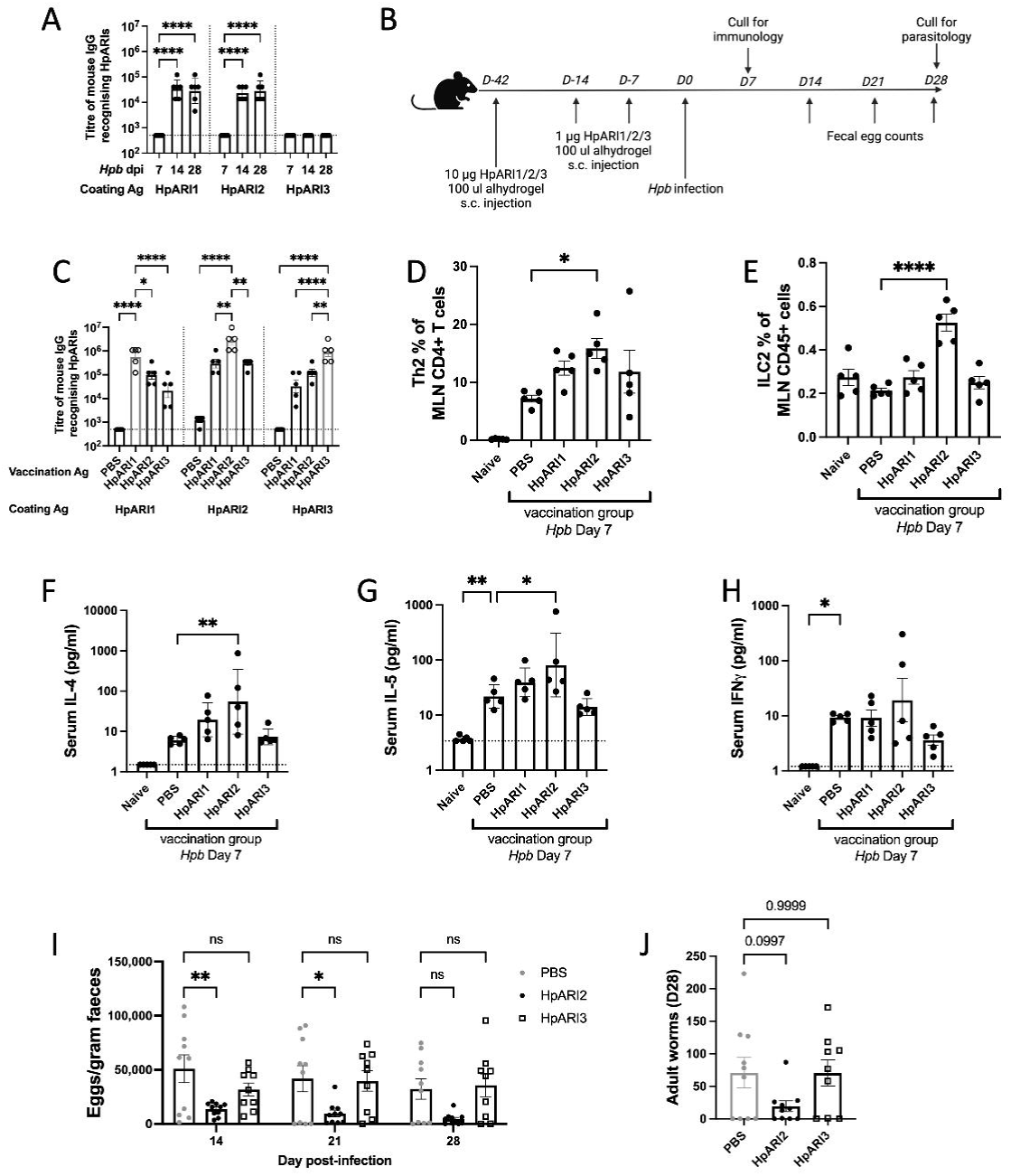
Vaccination with HpARI2 releases type 2 response from suppression and protects against *Hpb* infection. Mice were infected with *Hpb* and HpARI1, HpARI2 and HpARI3-specific serum IgG titres were measured at 7, 14 or 28 days post-infection (dpi) (A). Experimental protocol for vaccination experiments using HpARI1, HpARI2 or HpARI3 followed by *Hpb* infection (B). HpARI1, HpARI2 and HpARI3-specific serum IgG titres after vaccination, at day 7 of *Hpb* infection (C). Mesenteric lymph nodes were taken from vaccinated mice at day 7 of *Hpb* infection, to measure GATA3**^+^**Foxp3**^−^** percentage of mLN CD4^+^Lin^+^CD45^+^ Th cells (D), and GATA3^+^Lin^−^ ILC2 percentage of mLN CD45^+^ cells (E). Serum was taken from vaccinated mice at day 7 of *Hpb* infection, to measure concentrations of IL-4 (F), IL-5 (G) and IFN-γ (H). At days 14, 21 and 28 of *Hpb* infection after vaccination, *Hpb* eggs were counted in faecal samples (I). At day 28 of infection after vaccination, mice were culled and adult worm burdens were assessed (J). Dotted line on x axis indicates limit of detection, where relevant. Data representative of at least 2 repeat experiments, apart from HpARI1 group, which is from a single experiment. A, C-H and J were analysed by one-way ANOVA, I analysed by two-way ANOVA, all with Dunnett’s multiple comparisons test. In A, all groups were compared to day 7 of infection, in C all groups were compared to relevant vaccinated group (indicated in grey), in D-J all groups were compared to PBS vaccinated *Hpb* D7 control. A and C-H n=5 per group, I-J n=10 per group. Female C57BL/6J mice aged 6-8 weeks used in all experiments. Unless otherwise stated, differences are not significant. * = p<0.05, ** = p<0.01, *** = p<0.001, **** = p<0.0001

With the significant increase in type 2 immune responses on infection with HpARI2 vaccination, we further investigated the effect of vaccination of HpARI2 on parasite burden. In these experiments, we used HpARI3 as a control vaccine antigen, as it is a closely-related protein to HpARI2, however vaccination with HpARI3 showed no effects on type 2 immunity. HpARI2 vaccination resulted in the reduction in faecal egg counts from day 14 onwards (**Fig 2I**), with a trend for reduction in adult worm burden (**Fig 2J**). HpARI3 vaccination, by contrast, had no effect on parasite burden.

Together, this data indicates that HpARI2 vaccination results in a blocking antibody response against HpARI2, releasing the immune response from suppression, and allowing effective ejection of the parasite.

### HpBARI cocktail vaccination abrogates ST2 suppression

The same vaccination approach was taken to assess the role of the HpBARI family. Similarly to the response seen against the HpARI family, in natural infection we found no detectable antibody response against the HpBARIs at day 7 of infection (**Fig 3A**). While a significant antibody response was raised against HpBARI_Hom2, no response could be detected against HpBARI, which, like HpARI3, may reflect HpBARIs lower expression level compared to HpBARI_Hom2 ^28^. Vaccination with HpBARI and/or HpBARI_Hom2 produced a high-titre antibody responses against the relevant HpBARI, with some cross-reactivity between the two related proteins (**Fig 3B**).

**Figure 3:**
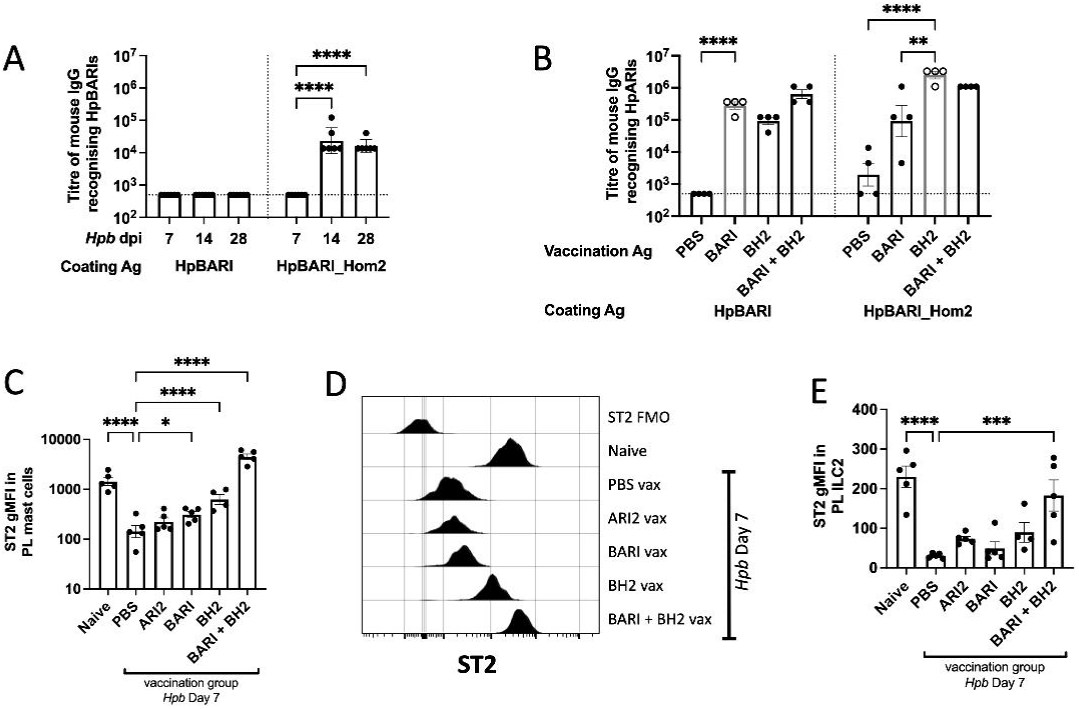
HpBARI family vaccination abrogates ST2 suppression. Mice were infected with *Hpb* and serum taken 7, 14 and 28 days post-infection (dpi) to measure HpBARI or HpBARI_Hom2-specific serum IgG titres (A). Mice were vaccinated with HpARI2 (ARI2), HpBARI (BARI) or HpBARI_Hom2 (BH2) as indicated, then infected with *Hpb* and culled at day 7 of infection. HpBARI (BARI) and HpBARI_Hom2 (BH2)-specific serum IgG responses (B). ST2 gMFI on peritoneal lavage (PL) CD45^+^Lin^−^ckit^+^FcERI^+^IgE^+^ mast cells (C) with representative histograms of ST2 on gated PL mast cells (D). ST2 gMFI on PL CD45^+^Lin^−^KLRG1^+^ ILC2 (E). Dotted line on x axis indicates limit of detection, where relevant. Analysed by one-way ANOVA with Dunnett’s multiple comparisons test, comparing all groups to PBS vaccinated *Hpb* D7 control. Unless otherwise indicated, differences are not significant. Data are representative of 2 repeat experiments. Female C57BL/6J mice aged 6-12 weeks used in all experiments. In A: n=6 per group, in B: n=4 per group, in C-E n=4-5 per group. * = p<0.05, ** = p<0.01, *** = p<0.001, **** = p<0.0001

To test the effects of blocking the HpBARIs, we first assessed the effect of vaccination on the abrogation of ST2 detection seen previously at day 7 of *Hpb* infection (as in **Fig 1A-C**). HpBARI family vaccination was compared to HpARI2 vaccination, as HpARI2 does not act on ST2 directly.

We found that vaccination with either HpBARI or HpBARI_Hom2 alone was not capable of fully reversing the ST2 suppression seen at day 7 of infection, but vaccination with both proteins in combination resulted in peritoneal cavity mast cells and ILC2 ST2 detection returning to (or exceeding) levels seen in uninfected mice (**Fig 3C-E**). HpARI2 vaccination, by contrast, did not significantly affect ST2 staining on these populations. Therefore, both HpBARI and HpBARI_Hom2 are required to fully suppress ST2 at day 7 of *Hpb* infection.

### HpBARI cocktail vaccination releases type 2 immunity from suppression and allows parasite ejection

We investigated the effects of HpBARI family vaccination in the mesenteric lymph node, finding that similarly to the peritoneal cavity, suppression of ST2 detection on ILC2 or Th2 was abrogated by combined HpBARI+HpBARI_Hom2 vaccination (**Fig 4A-B**). Surprisingly, in the mesenteric lymph node (but not in the peritoneal cavity), abrogation of ST2 suppression was achieved even with sole HpBARI_Hom2 vaccination (but not with sole HpBARI vaccination) indicating a site-specific difference in effects of these two immunomodulatory homologues. The combined HpBARI+HpBARI_Hom2 vaccination also increased type 2 immune responses, with increased proportions of Th2 cells (**Fig 4C**), but unlike HpARI2 vaccination (as in **Fig 2E**) there was no significant increase in ILC2 responses (**Fig 4D**). As with HpARI2 vaccination, increased type 2 immunity with HpBARI+HpBARI_Hom2 vaccination resulted in increased serum IL-4 and IL-5 (**Fig 4E-F**), but in contrast to HpARI2 vaccination, it also increased serum IFN-γ (**Fig 4G**). There was a smaller, but still significant effect of HpBARI_Hom2 vaccination alone, with no effect of HpBARI vaccination on these Th2 parameters. Altogether, these results indicate that to fully reverse the ST2 suppression seen in the early phases of *Hpb* infection, a cocktail of HpBARI+HpBARI_Hom2 is most effective.

**Figure 4:**
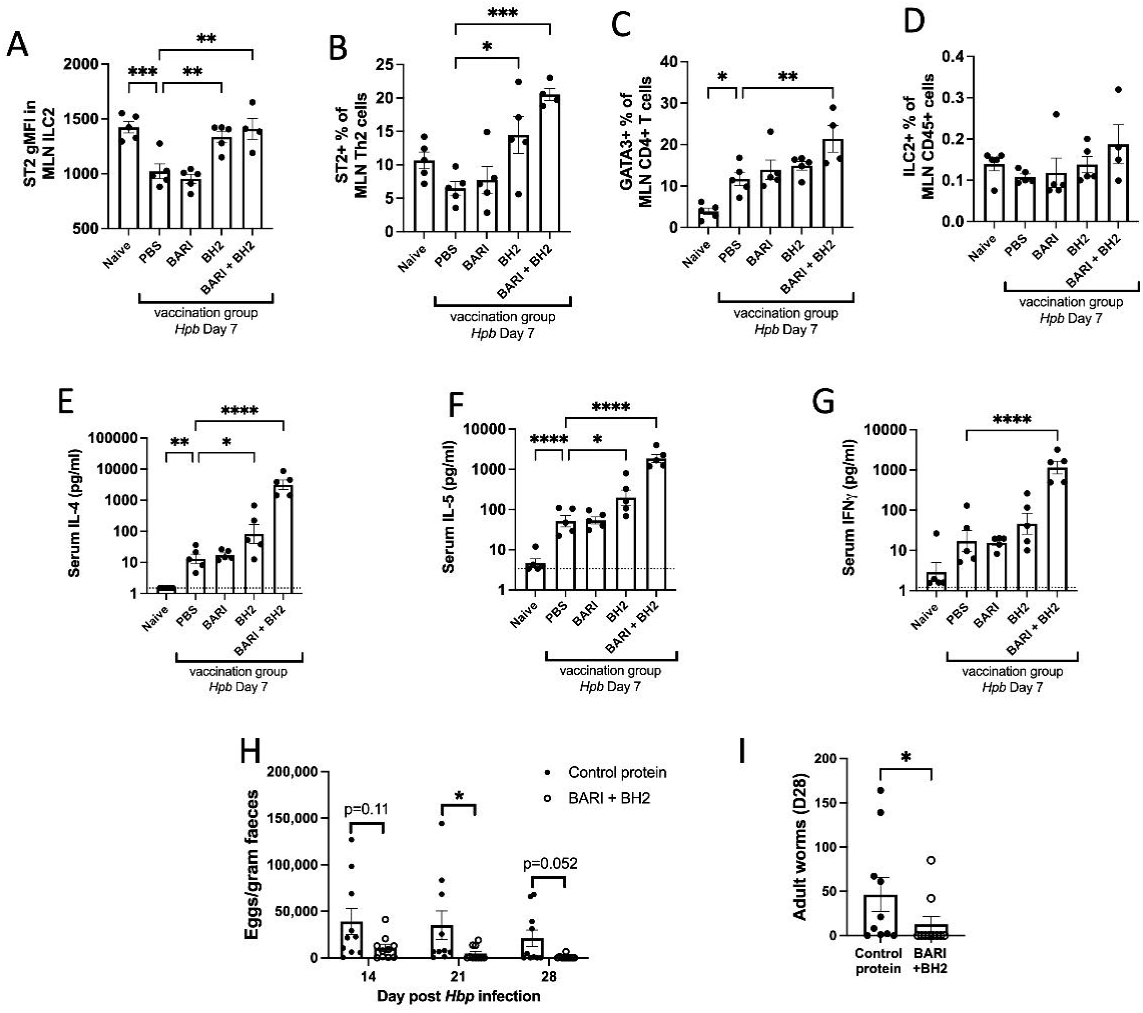
HpBARI family vaccination releases type 2 response from suppression. Mice were vaccinated with HpBARI (BARI) and/or HpBARI_Hom2 (BH2) and then infected with *Hpb*. At day 7 of infection, ST2 gMFI on MLN GATA3^+^CD127^+^Lin^−^CD45^+^ ILC2 (A), ST2^+^ % of MLN GATA3^+^CD4^+^Lin^+^CD45^+^ Th2 cells (B), GATA3^+^ % of MLN CD4^+^Lin^+^CD45^+^ Th cells (C), ILC2 % of MLN CD45+ cells (D). Serum IL-4 (E), IL-5 (F) and IFN-γ (G). Mice were vaccinated with HpBARI+HpBARI_Hom2 or HPOL_0001072601 control protein, and at day 14, 21 and 28 of *Hpb* faecal egg counts taken H). Vaccinated, infected mice were culled at day 28 of infection and adult worm burdens counted (I). Dotted line on x axis indicates limit of detection, where relevant. All data representative of 2 repeat experiments. A-G analysed by one-way ANOVA with Dunnett’s multiple comparisons test, H analysed by two-way ANOVA with Dunnett’s multiple comparisons test, while I was analysed by Mann-Whitney U test. In A-G, comparisons are made to PBS-vaccinated *Hpb* D7 control. In A-G, n=4-5 per group, in H-I n=10 per group. Female C57BL/6J mice aged 6-12 weeks used in all experiments. Unless otherwise stated, differences are not significant. * = p<0.05, ** = p<0.01, *** = p<0.001, **** = p<0.0001

To test the combination of HpBARI+HpBARI_Hom2, we vaccinated mice with a cocktail of these two proteins and used as a comparator another *Hpb* parasite-secreted protein expressed in the same expression system (HPOL_0001072601). HpBARI+HpBARI_Hom2 vaccination resulted in significantly reduced adult worm and egg burdens (**Fig 4H-I**). Together, these data show that the HpARI and HpBARI families have non-redundant roles in suppressing effective immune ejection of *Hpb*.

## Discussion

*Hpb* produces a range of immunomodulatory proteins which have been proposed to allow persistence of the parasite in vivo, preventing development of effective immunity. Here we show that vaccination with members of these immunomodulatory families was able to both induce clearance of the parasite, and abrogate the effects of these immunomodulatory proteins. This work therefore provides a proof-of-principle that these secreted proteins inhibit parasite ejection in a non-redundant manner, that immunomodulatory proteins have potential as vaccine candidates, and that vaccination can allow us to investigate the roles of these proteins in vivo.

An intractable problem in studies of parasitic helminths is the lack of suitable transgenesis systems. Here we show that vaccination allowed us to block specific parasite proteins in vivo, to assess their importance in parasite persistence. To our surprise, we found that members of the HpARI and HpBARI families which had similar effects in vitro had quite different effects when blocked in vivo. For instance, HpARI1 and HpARI2 similarly suppress IL-33 in vitro ^13, 14^, but only HpARI2 vaccination had a significant effect on the type 2 immune response during *Hpb* infection. Conversely, despite our recent findings that HpARI3 can stabilise IL-33 in vivo ^14^, vaccination had no effects on type 2 immunity during *Hpb* infection. In contrast, while HpBARI and HpBARI_Hom2 have very similar effects on suppression of ST2 in vitro ^15^, only vaccination with a combination of both proteins (and not each protein alone) fully abrogated ST2 suppression, and allowed active type 2 immunity to eject the parasite. Therefore our vaccination experiments indicate that the HpBARI family members act in a coordinated manner to block ST2, while vaccination with the HpARI or HpBARI families shows that these sets of proteins have non-redundant roles in infection.

As both the HpARI and HpBARI families act on the IL-33-ST2 pathway, we might expect that abrogating their effects through vaccination would result in identical outcomes. Indeed, vaccinating with either HpARI2 or HpBARI+HpBARI_Hom2 resulted in increased Th2 responses, increased IL-4 and IL-5 levels in the serum, and increased resistance to the parasite, indicating that the effects of both of these sets of proteins are required for suppression of the immune response and parasite persistence. However, some qualitative differences were identified between HpARI2 and HpBARI+HpBARI_Hom2 vaccinations: HpARI2 vaccination resulted in much increased ILC2 proportions in the draining lymph node (on which HpBARI+HpBARI_Hom2 vaccination had no effect), while HpBARI+HpBARI_Hom2 vaccination resulted in increased serum IFN-γ, which was not seen in HpARI2 vaccination. The reasons for the disparity between HpARI2 or HpBARI+HpBARI_Hom2 vaccination are unclear, but could be to non-classical IL-33 signalling. It was recently shown that oxidised human IL-33 was found to activate an (ST2-independent) RAGE-EGFR pathway ^29^. Our previous publications indicate that while HpARI2 binds only to reduced, and not oxidised IL-33, its binding may prevent the oxidation of the cytokine ^10, 12^, therefore HpARI2 may block both the ST2-dependent and RAGE-EGFR-dependent pathways. Meanwhile the HpBARI family’s effects appear constrained to ST2 ^15^, and would not affect oxidised IL-33 signalling.

Suppression of the IL-33 pathway by *Hpb* appears constrained to the early phases of infection, with suppression of IL-33-dependent responses in the Alternaria model and abrogation of ST2 detection only evident during the first week of infection. The transcription patterns of the HpARI and HpBARI family members correlate with this – while HpARI1 and HpARI3 remain relatively constant throughout the *Hpb* lifecycle, transcription of HpARI2 peaks in the first week of infection and is rapidly reduced thereafter ^14^. Similarly, while HpBARI transcription remains stable throughout infection, HpBARI_Hom2 also peaks in the first week of infection ^28^. Therefore, this first week of infection appears to be a critical window of immunomodulatory activity. During the first week of infection, *Hpb* resides within the wall of the duodenum, emerging into the lumen as adults at around day 7-10. Therefore, during this critical early time period, *Hpb* can release immunomodulatory factors directly into the tissue. The first week of *Hpb* infection is also associated with a weak type 1 response at the site of infection, which only switches to a type 2 response once the parasite enters the lumen ^30^. The action of the HpARI and HpBARI families may therefore inhibit the IL-33-mediated development of Th2 immunity until the parasite exits the tissue.

Although the HpARI and HpBARI families are clearly important to the immunomodulatory activity of *Hpb*, they are not the only immunomodulatory family secreted by the parasite. Many helminths, including *Hpb*, secrete active apyrases, and a cocktail vaccine consisting of 5 recombinant apyrases could provide partial protection from *Hpb* infection ^31^. Likewise, *Hpb* secretes the TGM family, which ligate the mammalian TGF-β receptor ^32, 33^. Although this family have been well characterised in vitro, their effects during infection are less well understood – vaccinating mice with TGM family members could allow dissection of their functions.

Our work indicates that parasite secreted immunomodulators can be used as vaccine antigens in helminth infection. Previous data in other parasite models supports this: vaccination with the *Trichuris muris* IL-13-inhibitory protein p43 provides protection against infection ^34^. In parasites of livestock, putative immunomodulatory proteins have been tested as vaccines, and also shown protection against infection ^35^. As there are currently no vaccines available for any human parasitic helminths, further investigation of immunomodulatory secreted products could lead to identification of effective vaccine candidates.

In summary, we provide proof-of-principle that vaccination with immunomodulatory parasite proteins can result in blocking antibody responses against these proteins, releasing the immune response from suppression, and allowing productive immunity to the parasite. This research highlights the importance of the identification of helminth immunomodulatory proteins for development as vaccine candidates, and provides a tool for investigation of immunomodulatory function during infection.

## Acknowledgements

This work was funded by a Wellcome Investigator award (221914/Z/20/Z) to H.J.M. We thank Prof Georgia Perona-Wright for feedback on the manuscript.

## Author contributions

D.J.S., S.H. and H.J.M. conceived and planned the study and wrote the manuscript. D.J.S., S.H., N.O., J.R., V.S. and H.J.M. conducted mouse experiments. T.F. conducted protein production and quality control. F.C. carried out in vitro IL-33 assays. D.J.S and T.F. maintained parasite lifecycle and parasitology measurements. All authors designed experiments, analysed data, and read and commented on the manuscript.

