## Supplementary Figures 1 to 5 for "Vaccination against helminth IL-33-modulators permits immune-mediated parasite ejection"

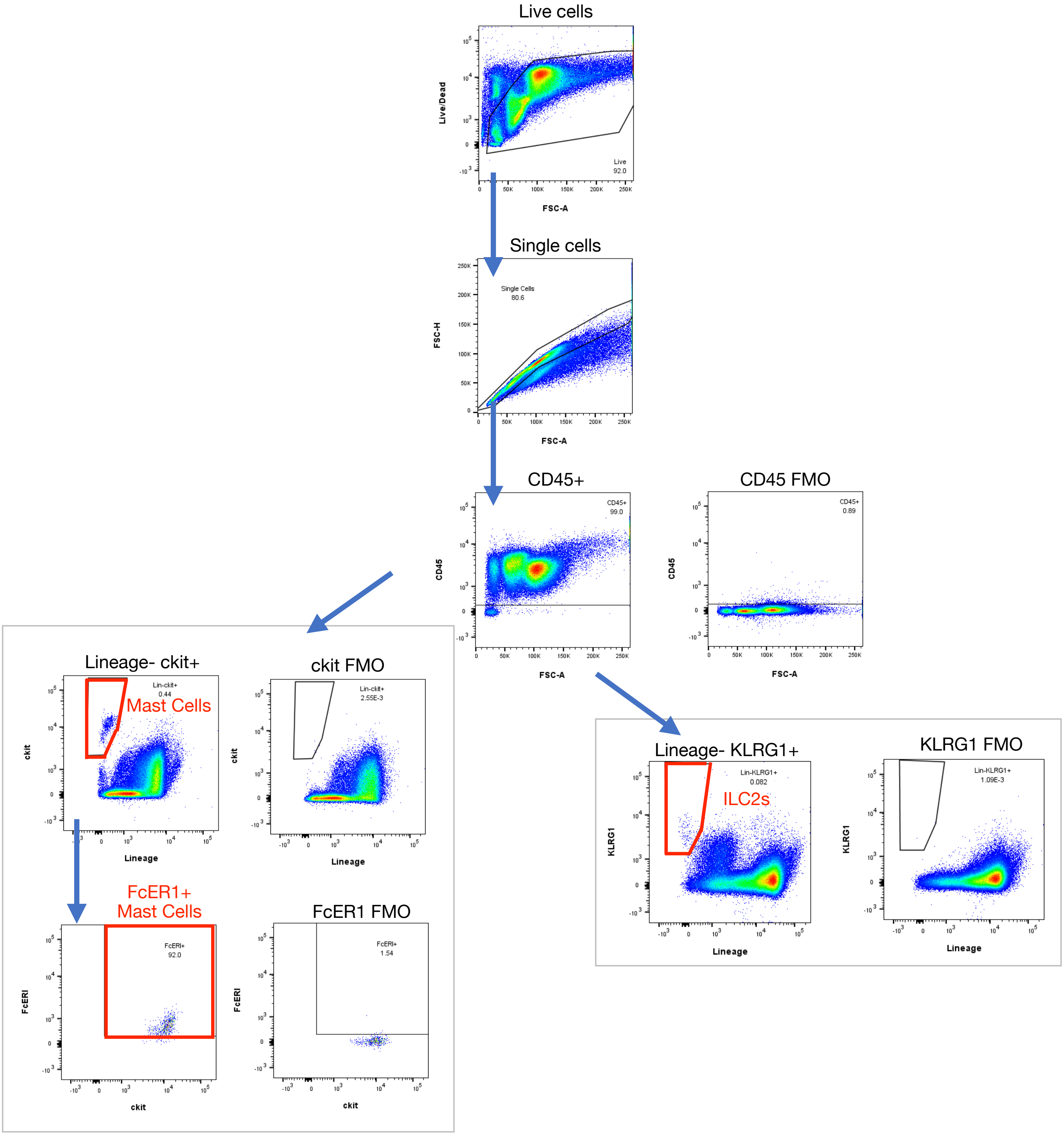


**Supplementary Figure 1:** Peritoneal lavage mast cell and ILC2 flow cytometry gating strategy

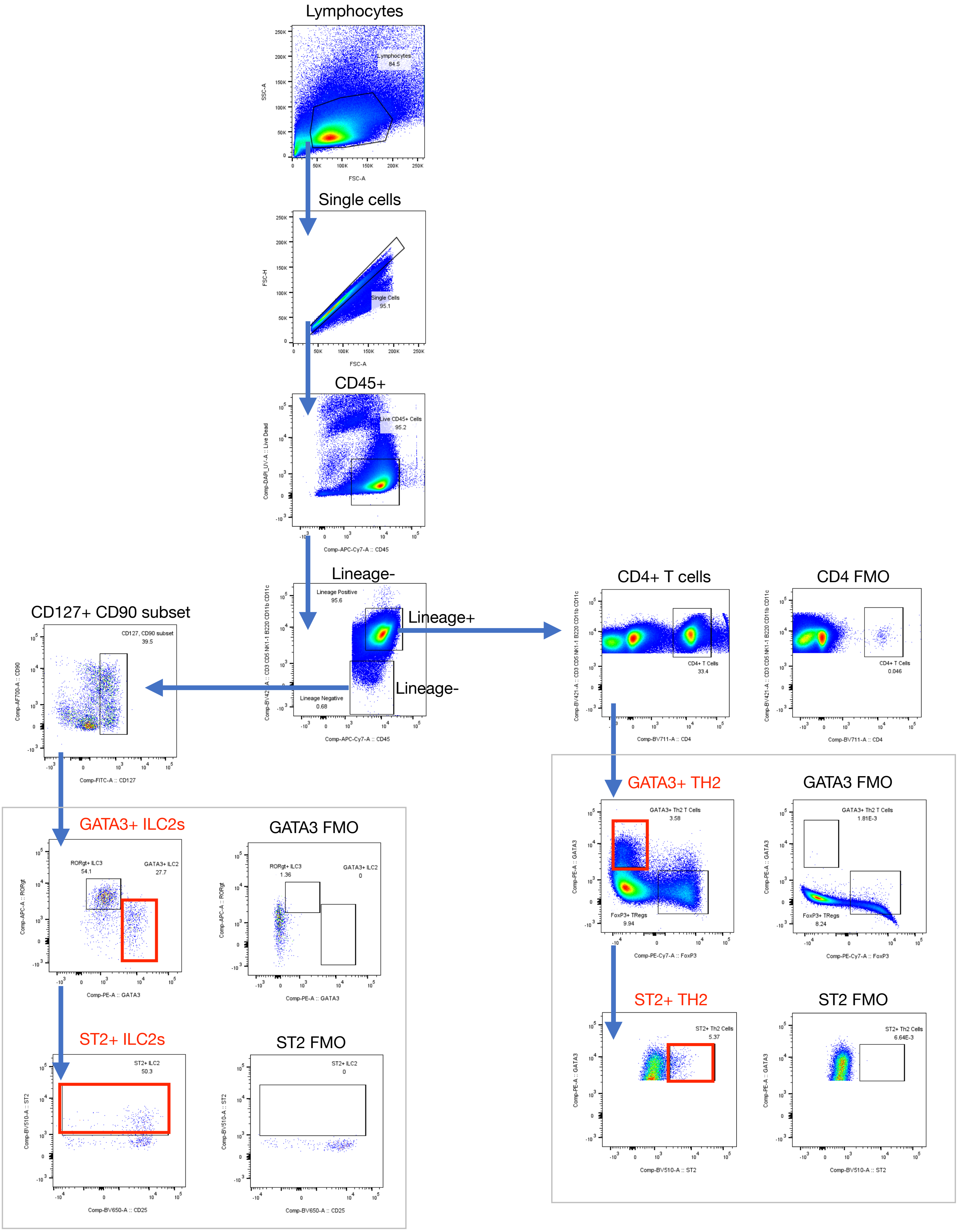


**Supplementary Figure 2:** Mesenteric lymph node ILC2 and CD4+ Th2 cell flow cytometry gating strategy

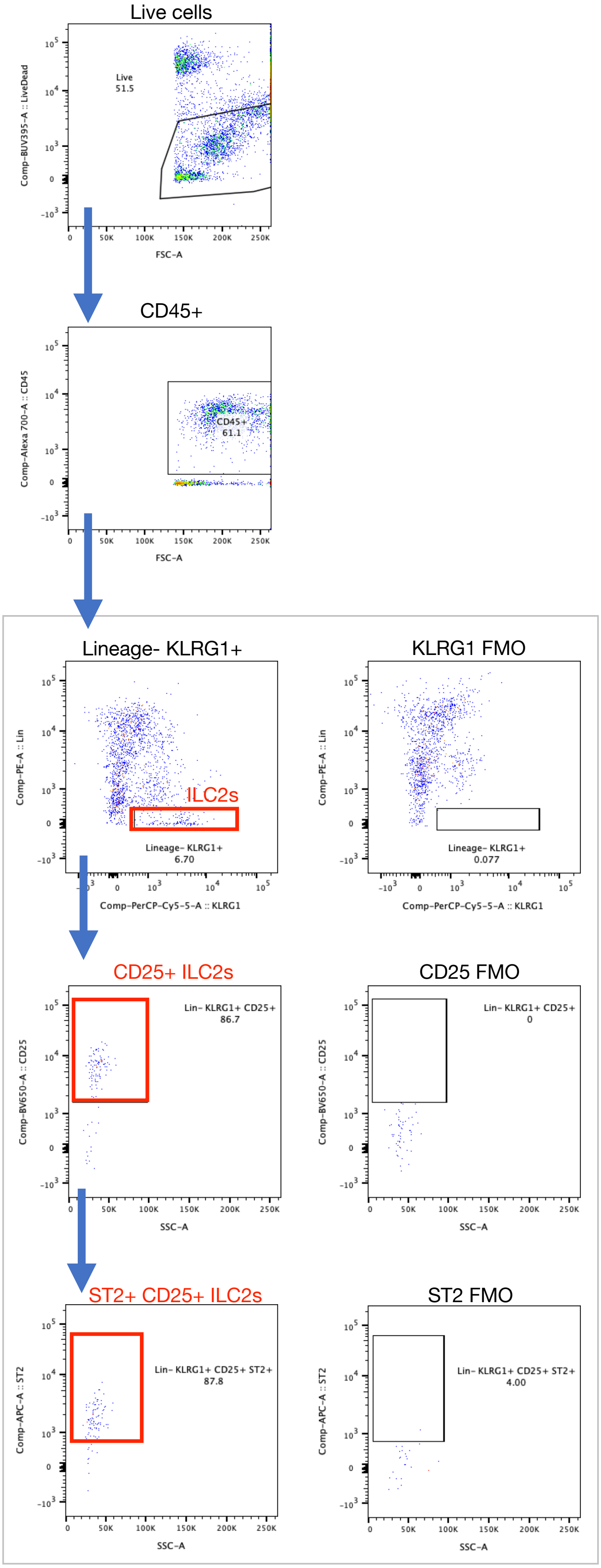


**Supplementary Figure 3:** pgWAT ILC2 flow cytometry gating strategy

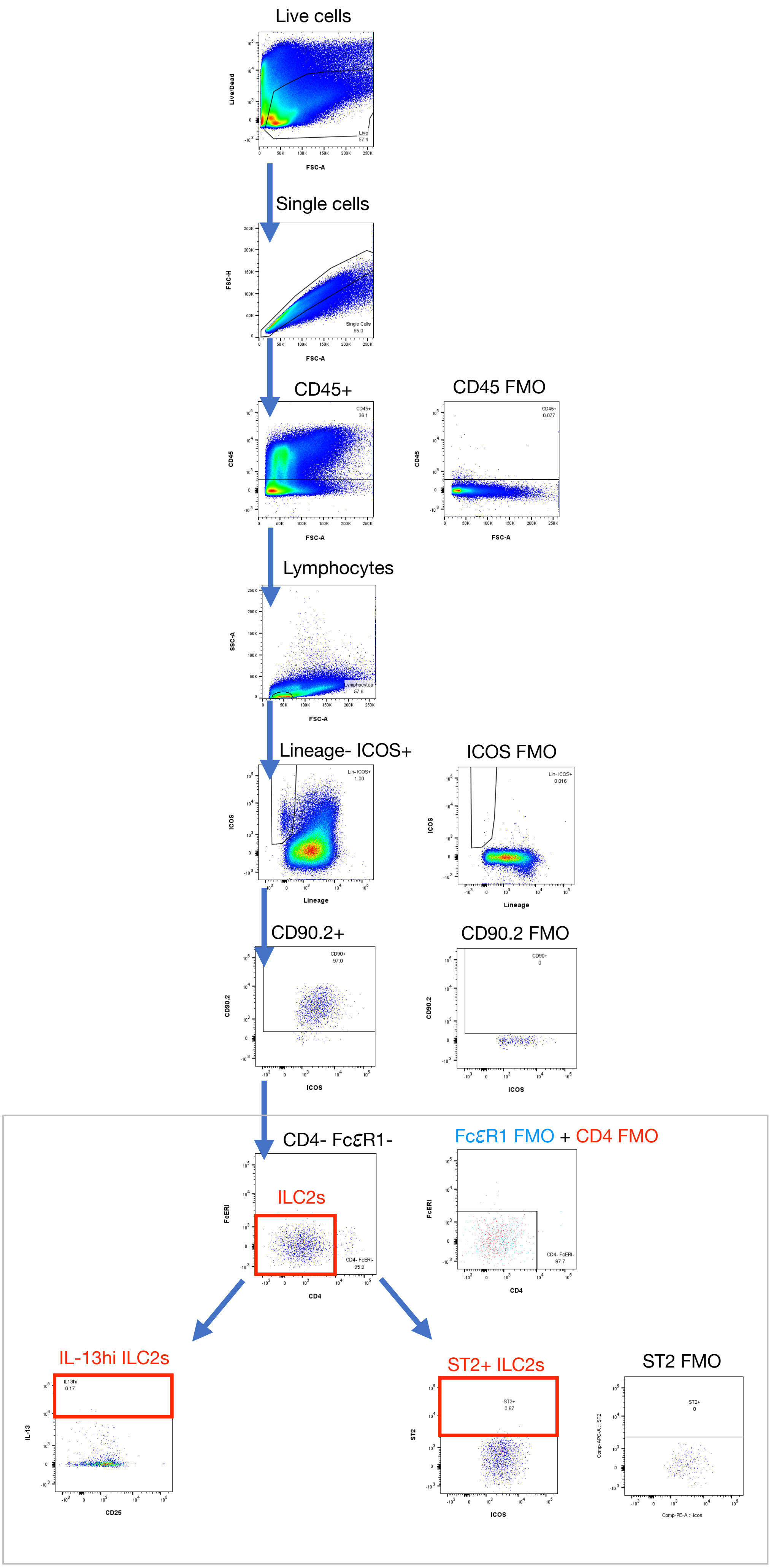


**Supplementary Figure 4:** Lung ILC2 flow cytometry gating strategy

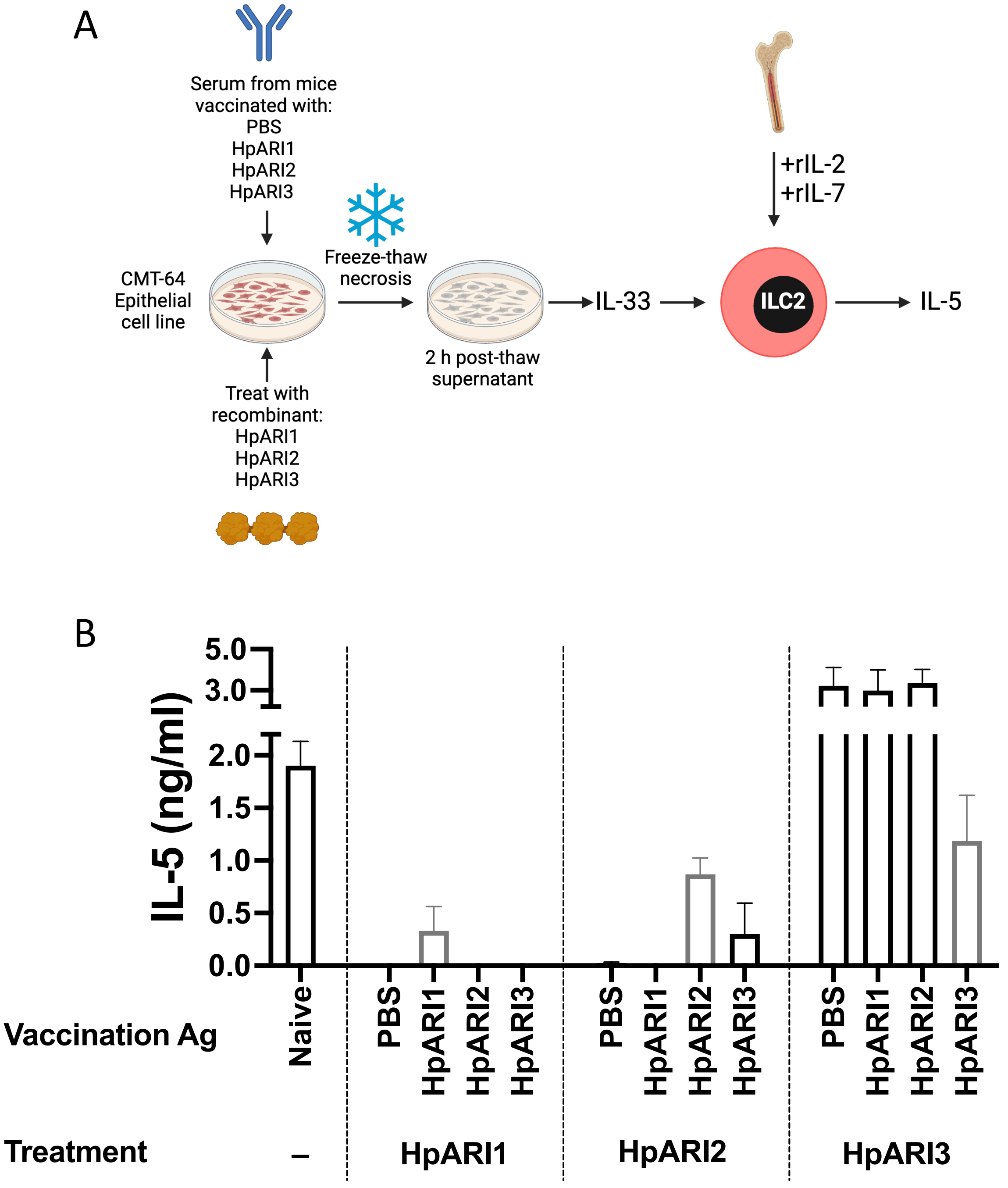


**Supplementary Figure 5: Vaccination serum to abrogate effects of HpARI1, HpARI2 or HpARI3 in vitro**

A. Experimental setup of IL-33 response assay. CMT-64 cells were freeze-thawed to induce necrosis and IL-33 release, in the presence or absence of HpARIs, and serum from mice at day 7 of *Hpb* infection, after vaccination with HpARIs. Supernatants containing IL-33 were taken from necrotic CMT-64 cultures 2 hours after thaw, and added to female mouse bone marrow cells in the presence of IL-2 and IL-7 (10 ng/ml each). Bone marrow cells were cultured for 5 days, and IL-5 in supernatants measured by ELISA.

B. IL-5 measurements in cultures as described in (A). Each group contains 5 replicates of serum, each from different vaccinated female mice (n=5).
